# Mechanically manipulate glymphatic transportation by ultrasound combined with microbubbles

**DOI:** 10.1101/2022.10.28.514316

**Authors:** Dezhuang Ye, Si Chen, Yajie Liu, Charlotte Weixel, Zhongtao Hu, Hong Chen

## Abstract

The glymphatic system is a perivascular fluid transport system for waste clearance. Glymphatic transportation is believed to be driven by the perivascular pumping effect generated by arterial wall pulsation induced by the cardiac cycle. Ultrasound sonication of circulating microbubbles in the cerebral vasculature induces volumetric expansion and contraction of microbubbles that push and pull on the vessel wall to generate a microbubble pumping effect. The objective of this study was to evaluate whether glymphatic transportation can be mechanically manipulated by focused ultrasound (FUS) sonication of microbubbles. The glymphatic pathway in intact mouse brains was studied using intranasal administration of fluorescently labeled albumin as a fluid tracer followed by FUS sonication at a deep brain target (thalamus) in the presence of intravenously injected microbubbles. Three-dimensional confocal microscopy imaging of optically cleared brain tissue revealed that FUS sonication enhanced the transport of fluorescently labeled albumin tracer in the perivascular space along microvessels, primarily the arterioles. We also obtained evidence of FUS-enhanced penetration of the albumin tracer from the perivascular space into the interstitial space. This study revealed that ultrasound combined with circulating microbubbles could noninvasively enhance glymphatic transportation in the brain.

**Significance Statement:** The glymphatic system is a waste clearance system in the brain analogous to the lymphatic system in peripheral organs. Glymphatic system impairment might contribute to brain disease pathologies, including those in neurodegenerative diseases, traumatic brain injury, and stroke. This study revealed that ultrasound could mechanically enhance glymphatic transportation. This result opens opportunities for using ultrasound to probe the role of the glymphatic system in brain function and brain diseases. Findings from this study suggest that ultrasound can be utilized as a noninvasive/nonpharmacological approach to mitigate brain diseases caused by impaired glymphatic function.

## Introduction

The glymphatic system is a glial-dependent perivascular network in the brain that functions in waste clearance and is analogous to the lymphatic system in the peripheral organs (1, 2). The glymphatic system was characterized for the first time in 2012 by Iliff *et al*. (3) in the mouse cortex. Iliff *et al*. showed that fluorescent tracers injected into the cerebral spinal fluid (CSF) entered the brain along cortical-penetrating arterioles through the perivascular space that was fully enwrapped by astrocytic endfeet. Their subsequent studies suggested that fluorescent tracers passed into the brain interstitium, moved toward the venous perivascular spaces, and ultimately moved into the cervical lymphatic system. The widely-accepted mechanism up to date for glymphatic transportation considers that glymphatic influx into and through the brain is driven, at least in part, by the perivascular pumping effect generated by arterial wall pulsation induced by the cardiac cycle (4). Indeed, carotid artery ligation significantly reduced arterial pulsatility and slowed the rate of perivascular exchange in the brain (5). Systemic administration of the adrenergic agonist dobutamine increased the pulsatility of penetrating arteries and enhanced CSF penetration into the parenchyma compared with the control (6). By contrast, angiotensin-II treatment induces hypertension and reduces the velocity of CSF flow (7).

The glymphatic system removes brain metabolic wastes such as excess protein and harmful metabolites in the interstitial space. The discovery of the glymphatic pathway provides new insights into how waste clearance works in the healthy brain, why we need to sleep, and the importance of efficient waste clearance in healthy aging (8). Impaired glymphatic transport is implicated in multiple neurological diseases, such as neurodegenerative diseases, traumatic brain injury, and stroke (8–14). Enhanced glymphatic transportation may mitigate brain diseases caused by impaired glymphatic clearance, and several strategies have been proposed to enhance glymphatic function. For example, mannitol treatment in mice led to plasma hyperosmolarity and a nearly 5-fold increase in CSF influx (15). Dietary supplements, such as omega-3 polyunsaturated fatty acids, promoted amyloid-β clearance in mice by promoting aquaporin-4 (AQP4)–dependent glymphatic transport (16, 17). Non-pharmacologic methods, such as sleep and physical exercise, have also been reported to upregulate glymphatic flow (18– 20).

We postulated that glymphatic transportation could be mechanically enhanced by ultrasound. Ultrasound, a noninvasive tool to generate mechanical effects, can penetrate the scalp and skull and reach the entire brain in animals and humans (21). Focused ultrasound (FUS), which concentrates ultrasound energy into a defined focal region, can target a specific brain area with millimeter precision (22). Effects from the FUS can be amplified at the targeted cerebral vasculature and surrounding brain tissue when combined with microbubbles (23). Microbubbles— compressible gas bubbles with a size comparable to red blood cells—were introduced in the clinic almost three decades ago as ultrasound contrast agents for imaging blood circulation (24). After intravenous injection, microbubbles are confined in the vasculature and circulate in the bloodstream like red blood cells. Ultrasound sonication leads to volumetric expansion and contraction of these compressible microbubbles, and these volumetric oscillations generate mechanical forces that push and pull on the adjacent vessel wall, thereby inducing vessel dilation and invagination (25). Vessel pulsation induced by FUS-activated microbubbles (FUSMB) generates the microbubble pumping effect.

Our previous studies reported that the microbubble pumping effect enhanced the penetration and accumulation of intranasally administered agents in the brain (26–30). Intranasally administered agents reach the brain through the olfactory and trigeminal nerve pathways (31). Once these agents reach the brain entry points, the perivascular pumping effect distributes them throughout the brain along the cerebral perivascular spaces (32, 33). Previous studies used invasive routes to study the glymphatic system (e.g., direct injection to the cisterna magna) (3). By contrast, intranasal administration provides a novel, noninvasive route to investigate the glymphatic pathway in intact brains. Our previous studies reported that FUSMB enhanced the accumulation of intranasally administered agents in the targeted mouse brain region (26–30); however, direct evidence is lacking to determine the effect of FUSMB on glymphatic transportation.

The objective of this study was to evaluate whether FUSMB can mechanically enhance glymphatic transportation. Fluorescently labeled albumin was intranasally administered to anesthetized mice, followed by FUS sonication of the mouse brain in the presence of intravenously injected microbubbles. Fluorescence imaging of the labeled tracer in optically cleared *ex-vivo* brain tissue revealed that FUSMB enhanced glymphatic transportation in the brain.

## Results

### FUSMB Enhanced Albumin Transport to the Targeted Brain Region

We applied FUS on the left side of the thalamus after intranasal administration of fluorescently labeled albumin (Fig. 1*A*). The contralateral right side was used as the control. Mice were sacrificed 15 min after sonication, their brains were harvested and sectioned into 1-mm thick coronal slices, which were then optically cleared for confocal fluorescence imaging (Fig. 1*B*). The acquired fluorescence images showed accumulation of the labeled albumin tracer in the FUS-treated thalamus region (Fig. 1*C*). Quantification of the fluorescence intensity of brain slices indicated that FUSMB enhanced the accumulation of albumin by 1.4-fold compared with the contralateral control. This result indicates that, at the macroscopic level, FUSMB enhances albumin transport to the FUS-treated brain region.

**Fig. 1.**
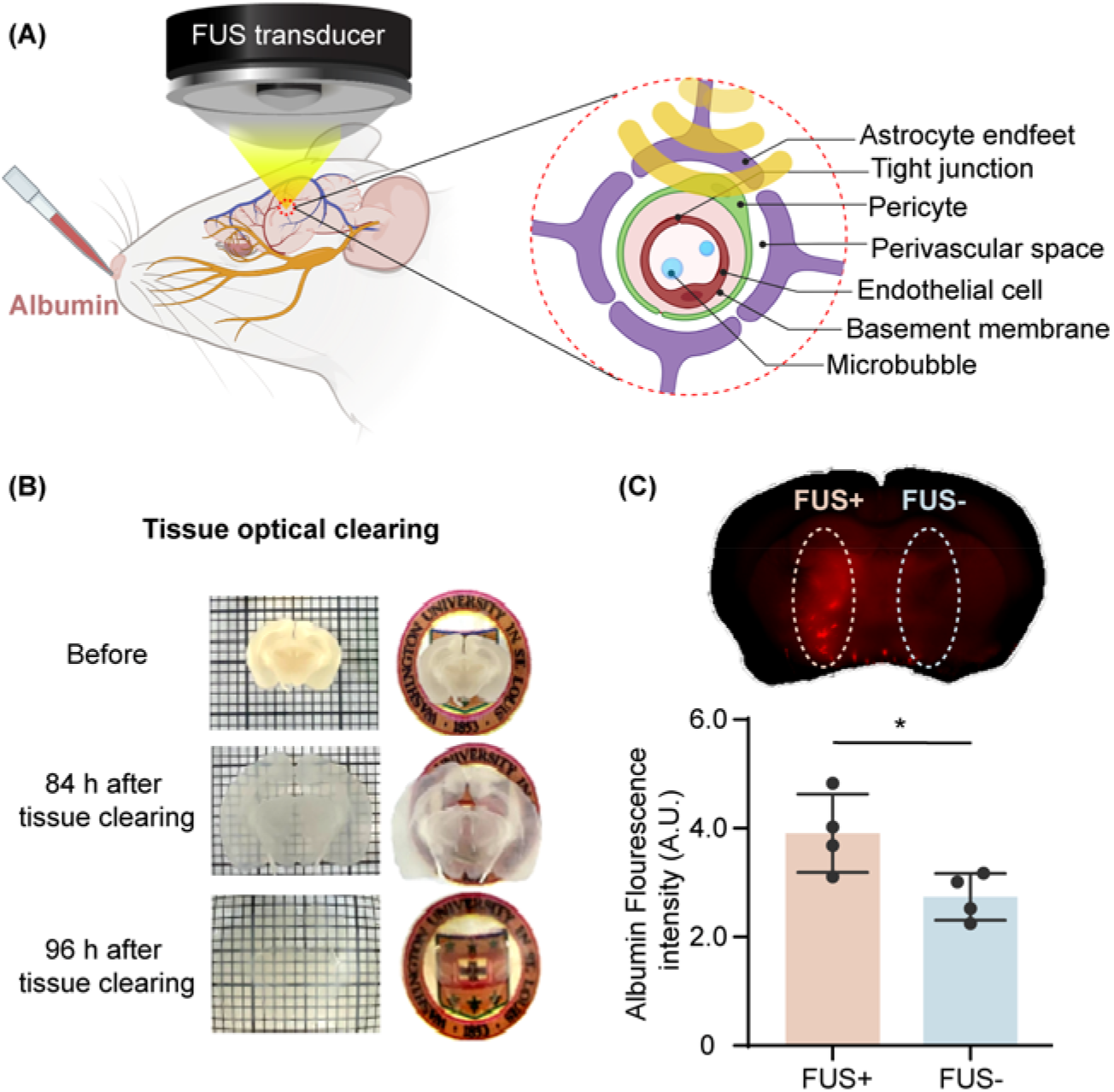
FUSMB enhanced albumin transport to the targeted brain region. (*A*) Illustration of the experimental method. Intranasal administration of fluorescently labeled albumin was followed by FUS sonication of a targeted brain region (left thalamus) after injection of microbubbles through the tail vein. (*B*) Harvested mouse brains were sectioned into 1-mm thick coronal sections and optically cleared using an established method. (*C*) Representative fluorescent image of albumin tracer in a 1-mm thick brain section (top). The fluorescence intensity of the albumin tracer in the FUS-treated side (FUS+) was significantly higher than that in the contralateral non-treated control side (FUS-) for a group of *N* = 4 mice (bottom). \**P* < 0.05.

### FUSMB Enhanced Albumin Transport via the Perivascular Space

To investigate the microscopic distribution of albumin following FUS, we stained optically cleared brain slices with lectin, a protein that binds to vessel walls, and anti-glial fibrillary acidic protein (GFAP) antibody, which identifies astrocytes. We found that albumin accumulated in the perivascular space between the vessel wall and astrocytic endfeet in the FUS-targeted brain region. This was consistently observed in both large (Fig. 2*A* and *SI Appendix* Movie S1) and small (Fig. 2*C* and *SI Appendix* Movie S2) vessels. By contrast, little to no albumin tracer was observed in the perivascular space on the contralateral side of the brain that did not undergo sonication (Fig. 2*B* and *D*). Cross-section images of the blood vessel confirmed that albumin is predominantly located in the perivascular space (Fig. 2*E* and *F, SI Appendix* Movie S3).

**Fig. 2.**
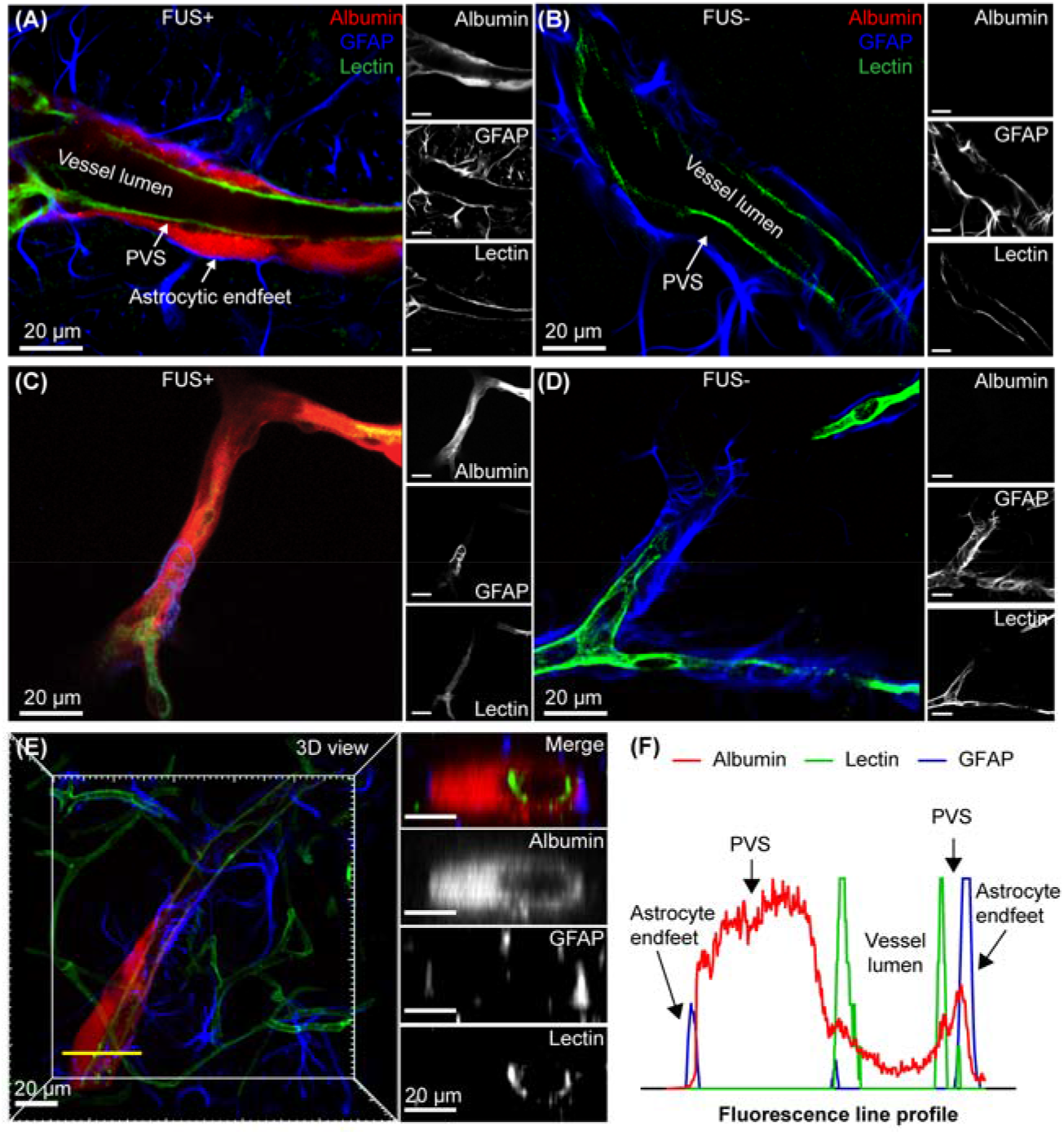
FUSMB enhanced albumin transportation along the perivascular space. Fluorescently labeled albumin (red) was observed in the perivascular space defined as the space between the lectin-stained vessel wall and the GFAP-labeled astrocyte endfeet of cerebral blood vessels in the FUS-treated brain region. (*A*) and (*C*) are representative confocal microscopy images of a relatively large blood vessel and a relatively small blood vessel in the FUS+ site, respectively. Little to no albumin tracer was observed in the contralateral non-treated (control) brain regions (*B* and *D*, respectively). (*B*) and (*D*) are representative confocal microscopy images of a relatively large blood vessel and a relatively small blood vessel in the FUS-site, respectively. (*A*–*D*) For each panel, the image on the left is a merged image of three channels (red, albumin; green, lectin; blue, GFAP) and the three images on the right are those obtained from individual images for albumin, GFAP, and lectin. (*E*) and (*F*) are representative cross-section images and the corresponding fluorescence profile, which verified that albumin was transported along the perivascular space. Images on the right panel of (*E*) show the cross-section images of a vessel captured along the yellow transect line in the 3D image shown on the left. The top image displays the merged image of three channels (red, albumin; green, lectin; blue, GFAP). The three images below show the individual images for albumin, GFAP, and lectin. Scale bars = 20 µm. (*F*) The fluorescence profile along the yellow transect line indicates that albumin accumulates in the perivascular space between the vessel wall and the astrocytic endfeet.

### FUSMB Enhanced Albumin Transport Primarily via Arterioles, Not Venules

To further investigate types of vessels that facilitate FUSMB-enhanced albumin transport, we stained cleared brain slices with anti-smooth muscle actin (αSMA) antibody and lectin to differentiate arteries from veins.. Three-dimensional (3D) confocal imaging of the FUSMB-targeted brain region showed that FUSMB significantly enhanced the transportation of intranasally administered albumin in the FUS-targeted brain region (Fig. 3*A* and *SI Appendix* Movie S4) compared with the contralateral brain (Fig. 3*B*). More importantly, we observed that albumin in the FUSMB-targeted brain region distributed primarily along the perivascular space of αSMA-positive vessels, i.e., arteries and arterioles (Fig. 3*A*). The corresponding *Z*-stack fluorescent images at different depths of the brain slice are presented in *SI Appendix* Figs. S1 and S2. To quantify the distribution of albumin along different types of vessels, we first computationally segmented individual vessels using the lectin staining. For each vessel, we measured the lumen diameter and quantified albumin localization by integrating the total albumin fluorescence intensity within 20 µm of the vessel surface (Fig. 3*C*). We identified arterioles, venules, or capillaries as described previously, according to smooth muscle cell coverage and vessel lumen size(34, 35). Consistent with our microscopy observation, the quantification of albumin distribution confirmed that arterioles had significant higher accumulation of albumin in its perivascular space compared to capillaries (p<0.05) and venules (p<0.01) (Fig. 3*D* and *E*).

**Fig. 3.**
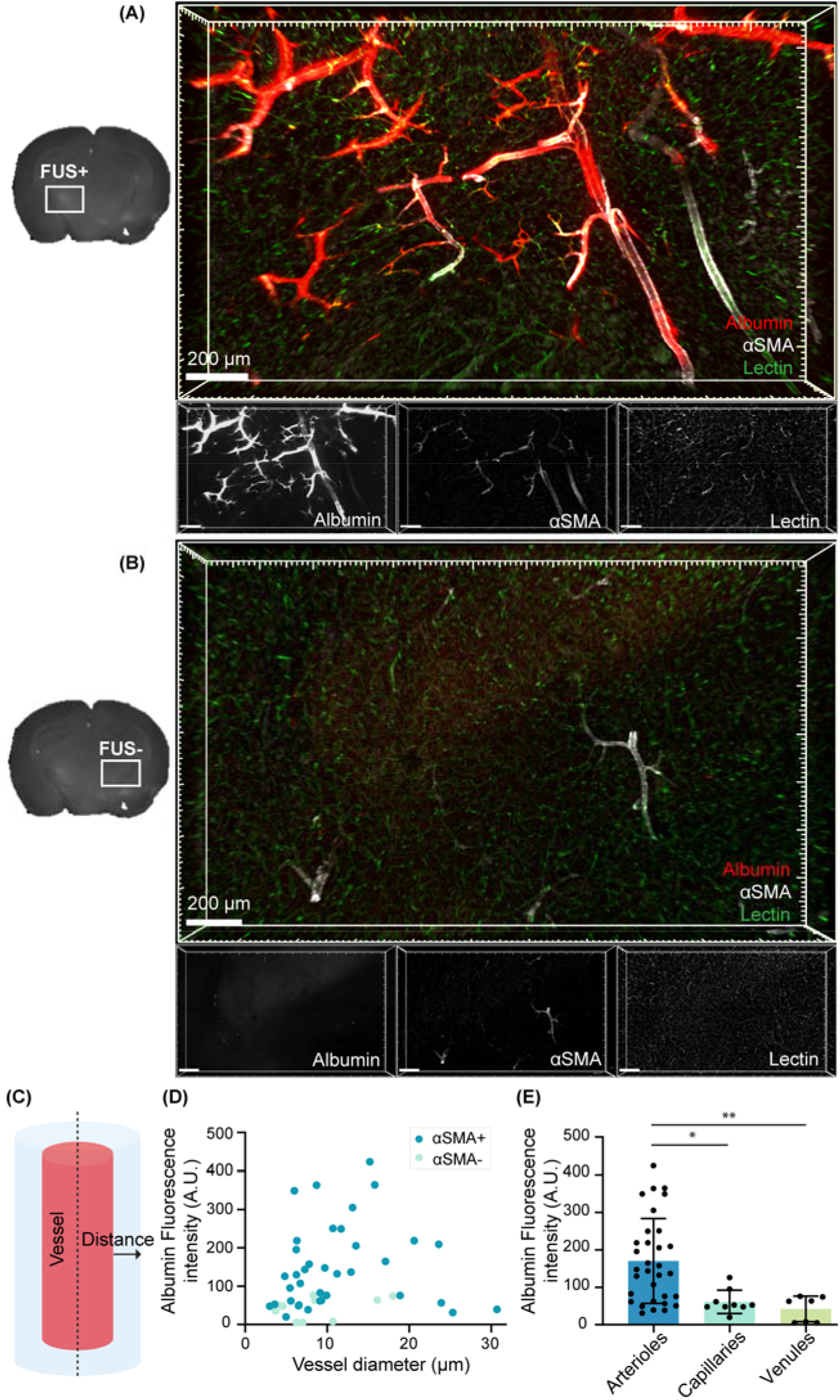
FUSMB enhanced albumin transportation primarily along arterioles, not venules. 3D confocal microscopy images were captured within a volume of 2×1.2×0.8 mm in (*A*) the FUS-treated brain region (FUS+) and (*B*) the contralateral non-treated (control) brain region (FUS-). For each panel, the image on the top is a merged image of three channels (red, albumin; green, lectin; white, αSMA), and the three images at the bottom display those obtained from individual channels. αSMA labeling was used to identify arterioles. Vessels with fluorescent albumin tracer accumulation (red) were either arterioles that were labeled with αSMA or capillaries that were connected to the arterioles. Scale bars = 200 µm. (*C*) Illustration of single vessel segmentation for quantifying albumin fluorescence intensity in the perivascular space. (*D*) Albumin fluorescence intensity of individual vessels in the FUS-treated brain region. Vessels that are αSMA+ and αSMA-are identified with different colors. (*E*) Albumin fluorescence intensity distribution for different vessel types. Arteriole: αSMA+, vessel lumen diameter >6 µm; capillary: vessel lumen diameter <6 µm; venule: αSMA-, vessel lumen diameter >6 µm. \**P* < 0.05; \*\**P* < 0.01.

### FUSMB Enhanced Albumin Transport into the Brain Interstitial Space

Confocal fluorescence images show that albumin can extravagate from the perivascular space of an arteriole into the interstitial space within 15 min after sonication (Fig. 4*A* and *SI Appendix* Movie S5). In addition to arterioles, we observed albumin in the perivascular space of capillaries and the surrounding interstitial area (Fig. 4*B* and *SI Appendix* Movie S6). Our findings suggest that FUSMB can enhance albumin penetration into the brain following extravasation from the perivascular space of arterioles and capillaries.

**Fig. 4.**
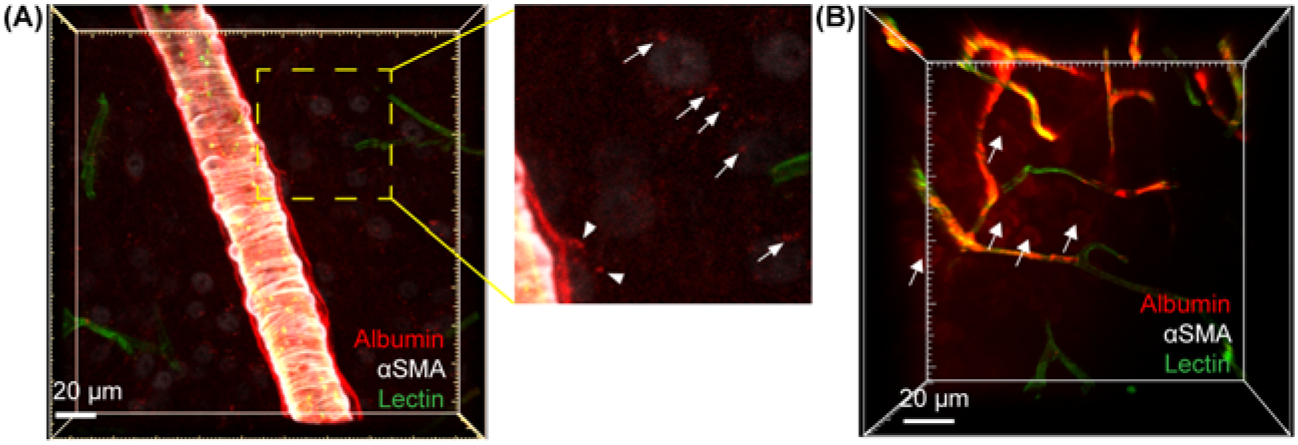
FUSMB enhanced albumin transportation into the interstitial space. Albumin extravasation from the perivascular space into the interstitial space was observed in arterioles (*A*) and capillaries (*B*) in the FUS-treated brain region. For (*A*), a zoom-in image within the yellow box is shown on the right, and white arrows mark albumin tracer in the interstitial space. For (*B*), the white arrows mark albumin tracer in the interstitial space. Scale bars = 20 µm.

## Discussion

The glymphatic system is a newly discovered brain fluid transportation system. This discovery presents enormous opportunities to develop strategies to investigate brain fluid dynamics and to treat brain diseases caused by impaired glymphatic transportation. Our study proposes that FUS combined with microbubbles can be used to mechanically manipulate glymphatic transportation in targeted brain regions, and that this enhanced transport is primarily through perivascular space of arterioles.

Our study obtained the first direct evidence that FUSMB enhanced the transport of a fluorescently labeled protein tracer in the perivascular space. Previously, Meng et al. observed that the MRI contrast agent accumulated in the draining veins and subarachnoid spaces following FUSMB in human brains (36). Lee et al. demonstrated that FUSMB enhanced soluble Aβ clearance from the brain to the CSF using a cervical lymph node ligation model in mice (37). These studies suggest that FUSMB may be used to study and manipulate the glymphatic system; however, the evidence presented did not show a direct effect of FUSMB on the glymphatic system. By contrast, our study using confocal microscopy imaging obtained evidence that unequivocally proved that FUSMB enhanced glymphatic transportation of a labeled protein agent in mice.

Our results show that FUSMB enhanced glymphatic transport primarily along arterials. The perivascular pumping effect generated by arterial wall pulsation has been considered the dominant driving force for glymphatic transportation into and throughout the brain. A previous study by Iliff et al. (3) reported that intracisternally injected tracer rapidly entered the brain along cortical surface arterials and penetrating arterioles. The tracer was not observed around veins at early time points (<10 min after injection). Similarly, our data show that FUSMB-enhanced albumin tracer accumulation occurred primarily in arterials, not venules, at 15 min after FUS sonication. Intravenously injected microbubbles circulate in the whole vascular network, including arterioles, venules, and capillaries. Our previous ultra-high-speed photomicrographic imaging of microbubble interactions with microvessels observed that FUS sonication–induced microbubble expansion and contraction generated pulsation-like behavior in arterioles, venules, and capillaries (38, 39). Thus, microbubble-induced vessel pulsation generates a microbubble pumping effect that may drive fluid transport along the perivascular space. The observed tracer distribution primarily in the perivascular space of arterioles supports the previous observation that glymphatic transport was driven by peri-arterial bulk flow and reveals that FUSMB enhanced the peri-arterial bulk flow. The observation of the albumin tracer outside the capillaries suggests that FUSMB may enhance directional flow in the perivascular space from arterioles to venules through the capillary.

Moreover, our findings showed that FUSMB enhanced the penetration of albumin from the perivascular space to the interstitial space. This is consistent with our previous reports that FUSMB increased the accumulation of intranasally administered agents (e.g., brain-derived neurotrophic factor, immune checkpoint inhibitor, and gold nanoparticles) in the FUS-treated brain region (27–30). Albumin was observed in the interstitial space 15 min post sonication, which is far more rapid than the speed of diffusion; therefore, this transport into the interstitial space is best explained by advection. Our previous ultra-high-speed photomicrographic imaging indicated that microbubble oscillations push and pull on the adjacent vessel wall and the surrounding connective tissue (38). Microbubble oscillations may generate a pressure gradient sufficient to drive directional flow from the perivascular space into the interstitial space. Thus, we suspect that the microbubble pumping effect contributed to agent transport along the perivascular space and penetration from the perivascular space to the interstitial space.

This study opens new opportunities to use FUS combined with microbubbles as a noninvasive and nonpharmacological approach to investigate and manipulate glymphatic transportation. Future studies are needed to quantify the distribution of different tracer agents at various time points and systemically characterize FUSMB effects on the glymphatic system. Further studies are also warranted to determine the correlations between different FUS parameters and the impacts on glymphatic transportation, which will optimize the FUSMB technique for glymphatic system investigations. It is critical to demonstrate that FUSMB enhances waste clearance in the brain and evaluate the potential therapeutic benefits of FUSMB in mitigating brain diseases caused by impairments in glymphatic system function.

## Materials and Methods

### Animals

All animal procedures were reviewed and approved by the Institutional Animal Care and Use Committee in accordance with the National Institutes of Health guidelines for animal research (IACUC protocol number 21-0187). Cr. National Institutes of Health (NIH) Swiss mice (6–8 weeks, ∼25 g body weight, female) were purchased from Charles River Laboratory (Wilmington, MA, USA). The animals were housed in a room maintained at 22°C, 55% relative humidity, and a 12-h/12-h light/dark cycle with access to standard laboratory chow and water until the experiment.

### FUSMB Procedure

Intranasal administration of Alexa Fluor 555-conjugated bovine serum albumin was performed as described previously (11). Briefly, mice were placed supine under anesthesia (1.5%–2% v/v isoflurane in oxygen). Eight drops (3 µL each) of 13 mg/mL fluorescently labeled albumin were administered to the mouse nose using a micropipette tip (Fig. 1*A*), alternating between the left and right nostril every 2 min. Following intranasal administration, mice were moved to a customized stereotaxic-guided FUS device to receive sonication under anesthesia as described previously (22). The fur on the mouse head was shaved, then ultrasound gel was applied to the exposed skin above the skull. A catheter was placed in the mouse tail vein for intravenous injection of microbubbles. A FUS transducer was positioned by the stereotaxic frame to target the left thalamus. Sonication started 30 min after intranasal administration of the labeled albumin. Immediately before sonication, mice were intravenously injected with a bolus of commercially available microbubbles (Definity^®^, Lantheus Medical Imaging, Billerica, MA, USA) at a dose of 10 µL/kg. The FUS parameters used in this study were as follows: pressure = 0.4 MPa, central frequency = 1.5 MHz, pulse length = 6.7 ms, pulse repetition frequency = 5 Hz, sonication duration = 60 sec.

### Tissue Clearing

Anesthetized mouse was transcardially perfused 15 min after FUS sonication using a perfusion pump at a speed of 5.5 mL/min. Blood of the mouse was first washed away by a 60 mL perfusion solution containing 1×PBS, 10 U/mL heparin, and 0.5% w/v sodium nitrite (Sigma Aldrich, St. Louis, MO, USA). The mouse was then fixed by transcardial perfusion with a 50 mL fixation solution containing 1×PBS and 4% w/v paraformaldehyde (PFA) (Sigma Aldrich). Mouse brain was harvested, and incubated in 4% PFA/PBS for 10 h at 4°C. The fixed brain was sectioned as 1-mm thick coronal slices using a brain matrix (RBM-2000C; ASI Instruments, Inc., Warren, MI, USA). Tissue clearance was performed as described previously (40). Briefly, slices were incubated in ScaleS0 solution for 12 h to permeabilize the samples. Then, the slices were incubated in ScaleA2 solution for 36 h, followed by incubation in ScaleB4 solution for 24 h, and re-incubated in ScaleA2 solution for 12 h. All incubations were performed at 37°C in an incubator shaker at 75 rpm (New Brunswick Scientific, USA). The samples were washed with 1×PBS at 4°C for 6 h. ScaleS0, ScaleA2, ScaleB4, and ScaleS4 were prepared using a previously published protocol (40).

### Macroscopic Fluorescence Imaging

After clearance, the brain sections were first imaged using an Olympus MVX10 microscope (Olympus, Japan). Images were captured using a color camera (Olympus DP23) at ×0.63 magnification, and images were saved using MetaMorph software (Olympus). The fluorescence intensities of the brain slices were quantified using ImageJ (NIH, Bethesda, MD, USA) by calculating the total fluorescence intensity of the FUS-treated side and the contralateral untreated side of the slices.

### Immunofluorescence Staining

Cleared brain slices were stained with αSMA to identify arterioles or GFAP to identify astrocytes. In brief, AbScale solution, AbScale rinse solution and ScaleS4 solution were prepared according to the previously published protocol (40). The primary antibody (αSMA [D4K9N, Cell Signaling Technology, Beverly, MA, USA] or Recombinant Anti-GFAP antibody [Abcam, ab207165, Cambridge, MA, USA]) was diluted 1:150 in AbScale solution. Then, brain samples were incubated in AbScale solution containing the primary antibody for 36 h at 37°C (with shaking). After incubation, the samples were washed twice in AbScale solution (2 h at room temperature with shaking) and then stained with the secondary antibody (Dylight 405 donkey anti-rabbit antibody [Antibody: AbScale solution, 711475152, Jackson Laboratories, West Grove, PA, USA]) for 48 h. Then, samples were washed with AbScale solution for 6 h and rinsed twice with AbScale rinse solution (2 h at room temperature with shaking). Samples were then refixed in 4% PFA/PBS for 1 h at room temperature with shaking and washed in 1×PBS for 1 h at room temperature with shaking. Samples were cleared in ScaleS4 solution for 12 h at 37°C with shaking and stored in ScaleS4 at 4°C until use for imaging.

### 3D Confocal Microscopy Imaging

Cleared and stained coronal brain sections were placed on a 35-mm glass-bottom dish (MatTek, Ashland, MA, USA). Images were acquired using a Zeiss Celldiscoverer 7 (Carl Zeiss, Toronto, ON, Canada). For the fluorescently labeled albumin tracer, an excitation wavelength of 561□nm and an emission wavelength of 575□nm was used for the channel setting; for the SMA and GFAP, an excitation wavelength of 405□nm and an emission wavelength of 421 nm was used for the channel setting; and for the FITC-labeled lectin, an excitation wavelength of 488□nm and an emission wavelength of 520 nm was used for the channel setting. Images within a large field of view were captured in the FUS-treated region and the contralateral non-treated region of the brain section using a 5× objective. The perivascular space was visualized in the FUS-treated region and the contralateral non-treated region of the brain section by capturing 3–5 images using a 20× objective.

### Single-Vessel Intensity Analysis

Single-vessel intensity analysis was performed using Imaris Version 9.8.0 (Oxford Instruments, Abingdon, UK). The Zeiss Celldiscoverer 7 images were imported, 3D reconstructed, and processed using the Imaris Gaussian filter. Background subtraction was applied to the 488 nm (lectin) channel. Then, a 3D surface was constructed from the 488 nm (lectin) channel to encompass all vessels in the sample. The lectin channel was masked by the resulting surface so that all background noise outside the vessel region was eliminated before single-vessel surface reconstruction. Next, the surfaces of individual vessels were computationally reconstructed for 12 vessels per subject on average. Each sample had approximately three images per region, which were analyzed separately or as one stitched image. Vessels of a variety of diameters were chosen from those displaying albumin tracer accumulation after intranasal delivery and FUSMB, thereby including a range of vessel diameters in the analysis. A surface was constructed to represent the spatial extent of each vessel based on lectin fluorescence. Vessel diameters were measured using the measurement function in Imaris.

### Statistical Analysis

Statistical analysis was performed using GraphPad Prism (version 8.3, La Jolla, CA, USA). Differences between the two groups were determined using an unpaired, two-tailed Student’s *t*-test. Differences among multiple groups were determined using one-way ANOVA followed by Bonferroni’s multiple comparison test. All groups passed the normality test. *P*-values < 0.05 were considered statistically significant.

## Author Contributions

D.Y., S.C., and H.C. designed the experiments. D.Y., S.C., and Y.L. performed the experiments. D.Y., S.C., Y.L., C.W., and Z.H. analyzed the data. D.Y., S.C., and H.C. wrote the paper.

## Data Sharing Plan

All study data are included in the article. Data that support the results of this study are available from the corresponding authors on reasonable request.

## Acknowledgments

This research was funded by the National Institutes of Health (NIH) grants R01EB027223, R01EB030102, and R01MH116981. The authors would also like to thank Dr. Joshua B. Rubin for his insightful comments and helpful discussion.

