## Supplementary information for "Mechanically manipulate glymphatic transportation by ultrasound combined with microbubbles"

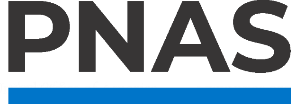

**Supplementary Information for**

**Mechanically manipulate glymphatic transportation by focused ultrasound combined with microbubbles**

Dezhuang Ye^1#^, Si Chen^1#^, Yajie Liu^1^, Charlotte Weixel^1^, Zhongtao Hu^1^, and Hong Chen^1,2*^

1. Department of Biomedical Engineering, Washington University in St. Louis, Saint Louis, MO 63130, USA.

2. Department of Radiation Oncology, Washington University School of Medicine, Saint Louis, MO 63130, USA.

\# These authors contributed equally.

* Address correspondence to: Hong Chen, Ph.D., Department of Biomedical Engineering and Radiation Oncology, Washington University in St. Louis, 4511 Forest Park Ave., St. Louis, MO, 63108, USA. Telephone: 314-454-7742.

**This PDF file includes:**

Figures S1 to S2

Legends for Movies S1 to S6

**Other supplementary materials for this manuscript include the following:**

Movies S1 to S6

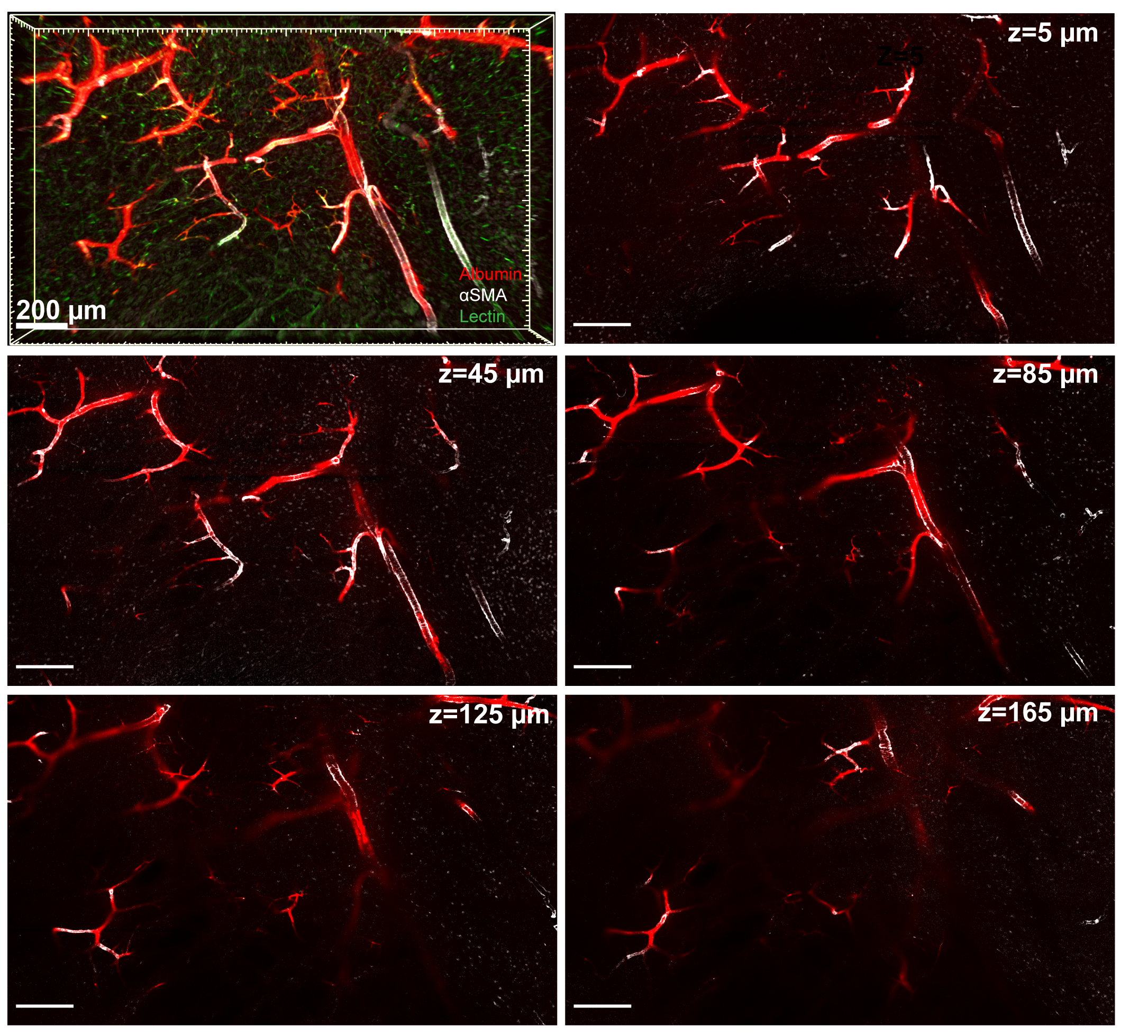

Fig. S1. Panel one shows the 3D image obtained from the FUS-treated brain region (also presented in Fig. 3A). The albumin fluorescence signal (red) is associated with the αSMA+ vessels (white). Other panels present individual images acquired at different depths with the depth (z) labeled in the upper right corner. Scar bar = 200 µm.

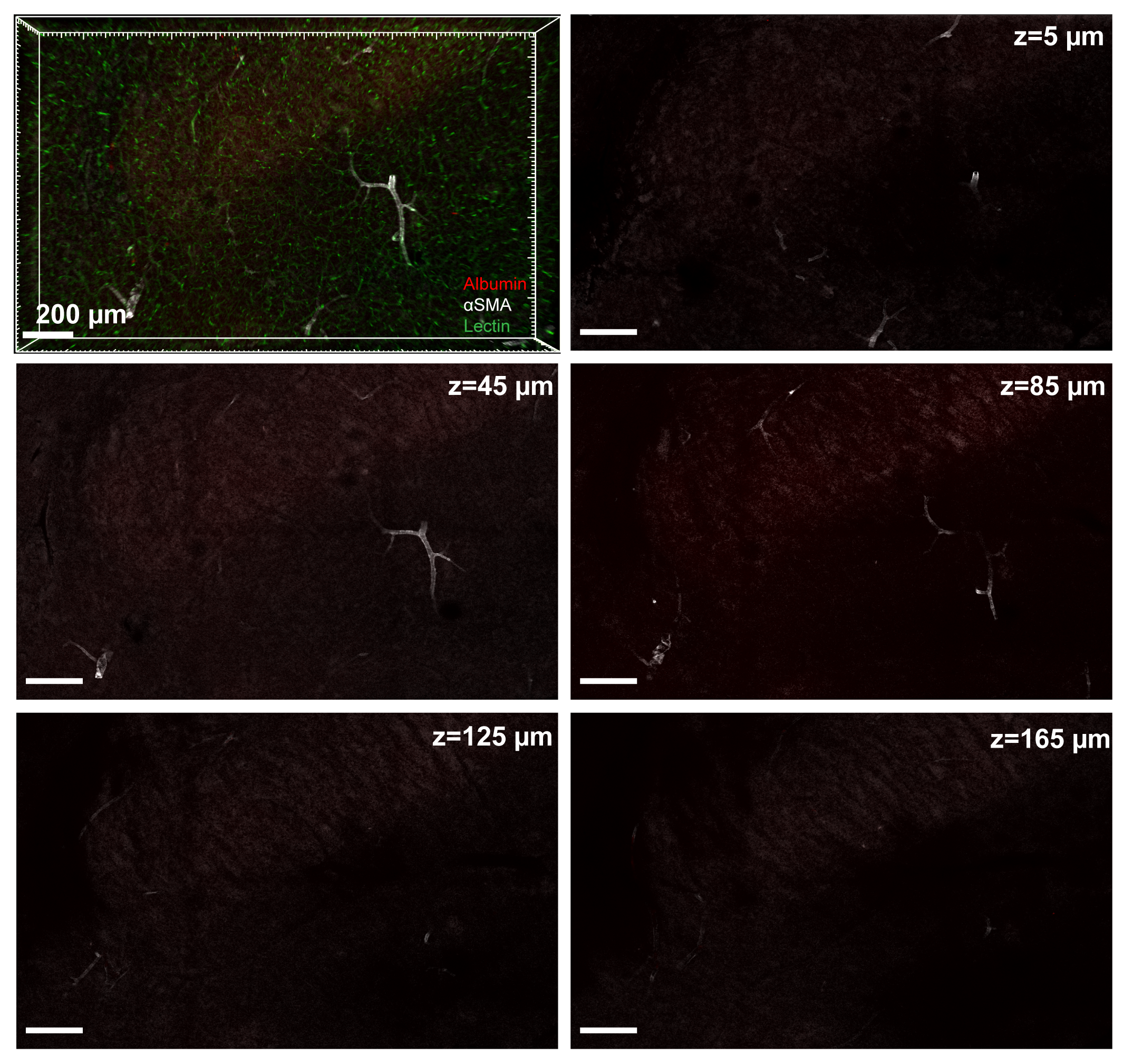

Fig. S2. Panel one shows the 3D image obtained from the contralateral non-treated control side (also presented in Fig. 3B). Other panels present individual images acquired at different depths with the depth (z) labeled in the upper right corner. Scar bar = 200 µm.

**Movie S1:** Three-dimensional view of the image shown in Fig. 2A with a voxel size of 0.139 μm × 0.139 μm × 2 μm. Red, albumin; green, lectin; blue, GFAP.

**Movie S2:** Three-dimensional view of the image shown in Fig. 2C with a voxel size of 0.139 μm × 0.139 μm × 0.46 μm. Red, albumin; green, lectin; blue, GFAP.

**Movie S3:** Three-dimensional view of the image shown in Fig. 2E with a voxel size of 0.139 μm × 0.139 μm × 2 μm. Red, albumin; green, lectin; blue, GFAP.

**Movie S4:** Z-stack view of the image shown in Fig. 3A. Red, albumin; green, lectin; white, αSMA.

**Movie S5:** Three-dimensional view of the image shown in Fig. 4A with a voxel size of 0.099 μm × 0.099 μm × 0.46 μm. Red, albumin; green, lectin; white, αSMA

**Movie S6:** Three-dimensional view of the image shown in Fig. 4B with a voxel size of 0.099 μm × 0.099 μm × 1 μm. Red, albumin; green, lectin; white, αSMA
